# Scaling principles of white matter brain connectivity

**DOI:** 10.1101/2021.05.31.445808

**Authors:** Dirk Jan Ardesch, Lianne H. Scholtens, Siemon C. de Lange, Lea Roumazeilles, Alexandre A. Khrapitchev, Todd M. Preuss, James K. Rilling, Rogier B. Mars, Martijn P. van den Heuvel

**Author notes:** Corresponding author, Dirk Jan Ardesch, De Boelelaan 1085, W&N B-636, 1081 HV, Amsterdam, the Netherlands. Email address.

## Abstract

Brains come in many shapes and sizes. Nature has endowed big-brained primate species like humans with a proportionally large cerebral cortex. White matter connectivity – the brain’s infrastructure for long-range communication – might not always scale at the same pace as the cortex. We investigated the consequences of this allometric scaling for white matter brain network connectivity. Structural T1 and diffusion MRI data were collated across fourteen primate species, describing a comprehensive 350-fold range in brain volume. We report volumetric scaling relationships that point towards a restriction in macroscale connectivity in larger brains. Building on previous findings, we show cortical surface to outpace white matter volume and the corpus callosum, suggesting the emergence of a white matter ‘bottleneck’ of lower levels of connectedness through the corpus callosum in larger brains. At the network level, we find a potential consequence of this bottleneck in shaping connectivity patterns, with homologous regions in the left and right hemisphere showing more divergent connectivity in larger brains. Our findings show conserved scaling relationships of major brain components and their consequence for macroscale brain circuitry, providing a comparative framework for expected connectivity architecture in larger brains such as the human brain.

## Introduction

Primate brains show a great diversity in shape and size, ranging from a few cubic centimeters in mouse lemurs and galagos to around 1300-1400 cm^3^ in humans (Stephan et al. 1981; Schoenemann 2013). From smaller to larger brains, however, not all structures tend to keep similar proportions. Post-mortem and early MRI studies across primate species have suggested that some structures grow faster than others, a phenomenon known as allometric scaling (Hofman 1989; Rilling and Insel 1999a). For example, the cerebral cortex scales faster than expected based on total brain volume, resulting in a proportionally larger cortex in humans and other great apes compared to smaller primate species (Hofman 1989; Mota et al. 2019). The effect of allometric scaling on the network structure of macroscale brain connectivity, a key factor in shaping brain function (Sporns et al. 2005), remains largely unknown. How do brain scaling principles affect these wiring patterns in the brain?

The anatomical substrate of region-to-region connectivity in the mammalian brain is the white matter, which has been noted to take up more and more space in larger-sized brains compared to smaller brains (Deacon 1990; Rilling and Insel 1999a; Hofman 2001). White matter accounts for 11% of total brain volume in mice and increases to 27% in macaques and more than 40% in humans (Zhang and Sejnowski 2000; van den Heuvel et al. 2016). While this may be the case, the total surface area of the cortex has been found to scale even faster (Hofman 1989; Mota et al. 2019), which has led to the theoretical question of whether the increasing white matter proportion can actually match the demand of interregional cortical connectivity in an expanding brain. The rapidly increasing number of cortical neurons has been theorized to outpace the space available for axonal connections between these neurons in different areas (Ringo 1991; Herculano-Houzel et al. 2007), suggesting a net decrease in direct corticocortical connectivity in larger brains. Evidence for this theory can be found at the neuronal level, where larger brains have been reported to display a lower fraction of cortical neurons that project directly into the white matter (Herculano-Houzel et al. 2010), placing strong constraints on ‘expensive’ long-range white matter connectivity between brain areas. Increases in brain size may thus impact overall brain connectedness, especially through long-range projections, and may similarly have effects on the global organization of brain connectivity.

To examine the effects of scaling on the organization of macroscale brain connectivity we compared brains of fourteen primate species across three orders of magnitude in size. We argue that a constraint on long-range connectivity may have the strongest effect on costly long-range white matter projections like the corpus callosum, the main body of fiber bundles connecting areas across the two hemispheres. We thus hypothesized that with relatively less space for callosal fiber bundles, interhemispheric connectivity could be particularly constrained, forming a ‘bottleneck’ that might prompt higher levels of hemispheric lateralization in larger brains. We reconstructed whole-brain connectivity maps and examined allometric scaling relationships between volumetric and connectivity measures. We first validated the findings of early post-mortem and MRI studies (Stephan et al. 1981; Rilling and Insel 1999a; Mota et al. 2019) showing that although white matter outpaces gray matter, this increasing proportion of white matter cannot keep up with the faster growth of cortical surface area in the primate brain. Second, we then zoomed in on the white matter of the corpus callosum to examine the impact of brain size on the space available for interhemispheric connectivity. Finally, we investigated the implications of scaling principles on network organization of the brain, showing that connectivity patterns between cortical regions in the left and right hemispheres become more and more asymmetrical with increasing brain size. Taken together, our findings suggest that brain scaling principles extend to patterns of macroscale brain connectivity and support the notion of growing levels of lateralization of brain structure in big-brained species, like humans.

## Materials and Methods

### MRI data

MRI data were included from a total of fourteen species (Figure 1). These include all extant great ape species (human, chimpanzee, bonobo, gorilla, orangutan), an ape species (lar gibbon), three Old World monkey species (rhesus macaque, grey-cheeked mangabey, black-and-white colobus), four New World monkey species (tufted capuchin, night monkey, woolly monkey, white-faced saki), and a prosimian (Senegal galago). MRI acquisition protocols, demographics, and data sources are detailed in Table 1. Data were included from an ongoing cohort of MRI scans of brains of the Primate Brain Bank (primatebrainbank.org/digital; technical details described in Bryant et al., in press), the National Chimpanzee Brain Resource (chimpanzeebrain.org) and data from previous studies (Rilling and Insel 1999a; Allman et al. 2010; Li et al. 2013; Cabeen et al. 2020). All in-vivo data in the National Chimpanzee Brain Resource were acquired prior to the 2015 implementation of US Fish and Wildlife Service and National Institutes of Health regulations governing research with non-human primates. Human procedures were approved by Emory University’s Institutional Review Board (IRB00000028), and all participants provided voluntary informed consent. We aimed to maximize the number of species that could be included by collating data from both in vivo and post-mortem samples, an approach that has been shown feasible in previous comparative tractography studies (Roumazeilles et al. 2020).

**Figure 1:**
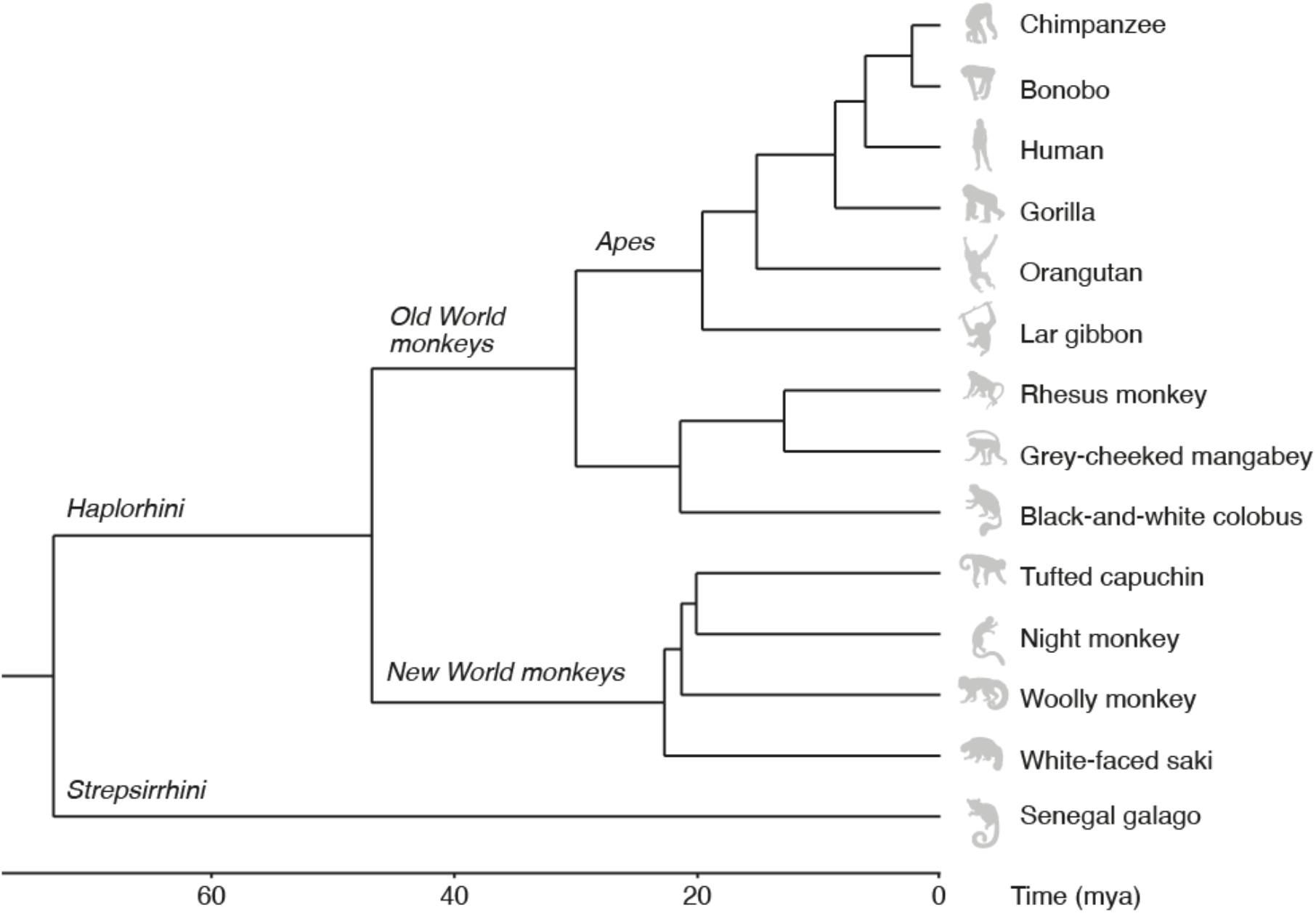
Phylogram of divergence times in million years ago (mya) for the primate species included in this study. Phylogeny was estimated from a consensus tree based on genotyping data of seventeen genes using version 3 of the 10kTrees project (Arnold et al. 2010). Images from phylopic.org.

**Table 1:**
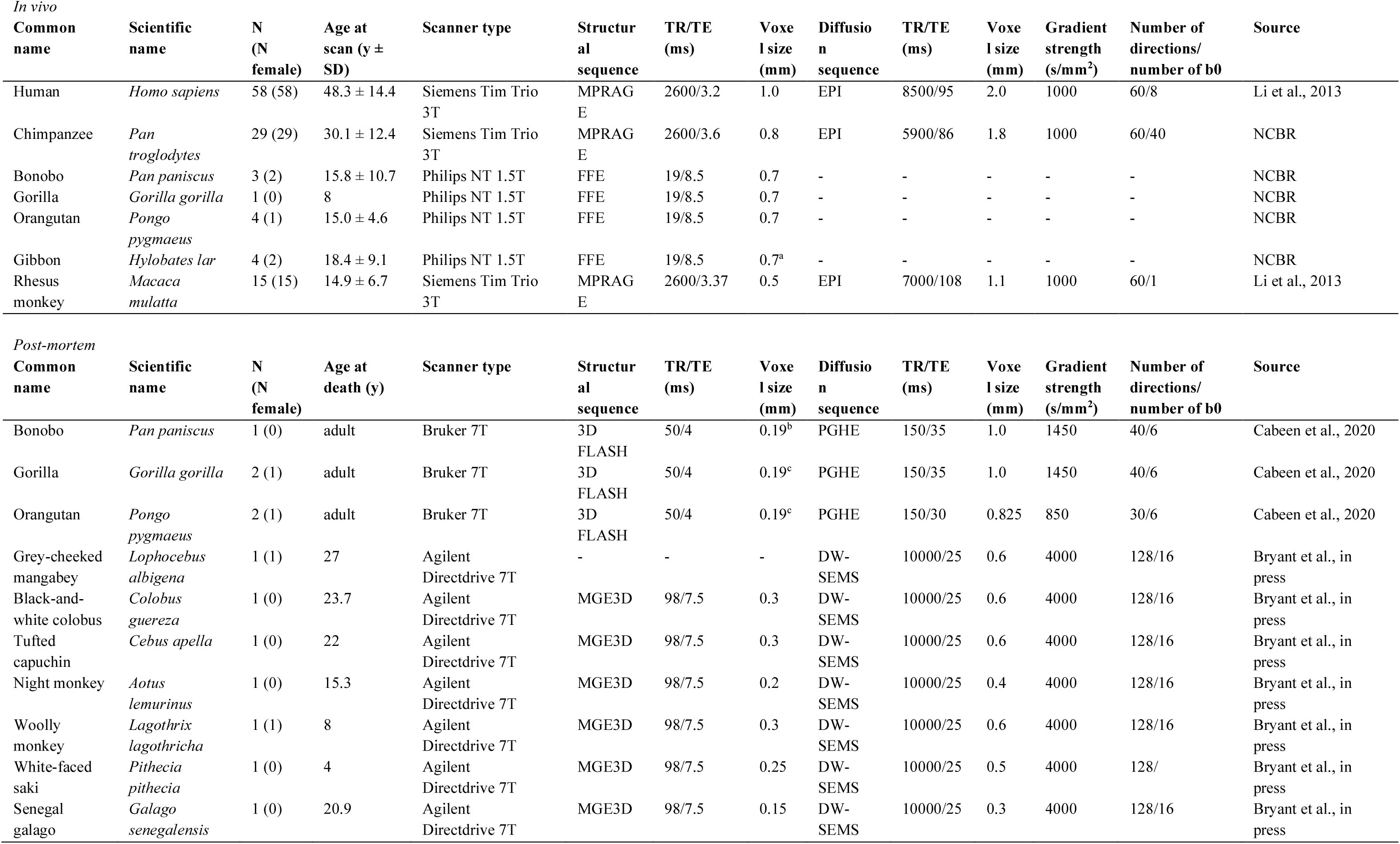
Demographics, MRI scan parameters, and sources of the datasets used in this study. If subjects within a dataset were acquired with different sets of parameters, the most common set of scan parameters is reported. Voxel sizes are isotropic unless noted by a superscript letter: ^a^0.7×0.7×0.6; ^b^0.19×0.22×0.29; ^c^0.19×0.22×0.33.

### Structural MRI processing

Structural MRI scans were processed using the FreeSurfer v6.0 pipeline (Fischl 2012), including tissue segmentation of cortical gray and white matter, subcortical structures, and reconstruction of cortical surfaces. For non-human primate datasets the FreeSurfer pipeline was complemented with tools from FSL v6.0.1 (Jenkinson et al. 2012), ANTs (Avants et al. 2011) and MATLAB to obtain surface reconstructions. Cortical reconstructions were visually inspected for accuracy and consistency across datasets. A brain mask was used to separate brain tissue from any other structures present in the scans such as skull (in vivo samples) or fiducial markers (post-mortem samples). Bias field correction was then applied to the images using ANTs. Voxel intensities of T2*-weighted images (see Table 1 for datasets) were inverted to create a contrast with low intensity for gray matter voxels and high intensity for white matter voxels. Tissue segmentation was performed using FreeSurfer (Figure S1). Manual corrections were made where needed to correct segmentations of white matter, cerebellum, and subcortical structures. FreeSurfer’s default Talairach registration was complemented by a step-by-step registration process for the smallest brain samples, starting with a registration from the subject at hand to a macaque template brain (Seidlitz et al. 2018), followed by a registration from the macaque template to a chimpanzee template brain (based on the chimpanzee subjects in this study), and from the chimpanzee template to human Talairach space. This step-by-step registration process ensured that major brain structures such as the cerebellum and thalamus were correctly segmented.

### Volume and surface metrics

Volume estimates of cerebrum (i.e., total brain volume except cerebellum and brainstem), cerebral white matter, and cortical gray matter were extracted from the FreeSurfer segmentation and statistics files of the examined brains. The same files were used to compute total cortical surface area and average cortical thickness. The corpus callosum (CC), a major interhemispheric white matter tract in primates, was clearly identifiable in all examined species so we included its cross-sectional area (i.e., the maximum area of the corpus callosum segmentation in the sagittal plane) as a measure of CC size. Surface area of the midsagittal slice of the CC is a good estimate of total interhemispheric connectivity (Elahi et al. 2015) with fibers running in a consistent direction perpendicular to the sagittal plane.

### Cortical parcellation

The cortical mantle was divided into a set of distinct, randomly placed areas (Arslan et al. 2018). A random parcellation was used to accommodate the lack of a biologically-informed atlas that maps the same, homologous regions in all of the examined primate species. The parcellation procedure consisted of placing a fixed number of region centers evenly dispersed across the cortical mantle. Each vertex in the cortical surface reconstruction was then assigned to the closest region center, resulting in a fixed number of regions of approximately equal size (50 regions per hemisphere, analyses using 25 or 100 regions per hemisphere are described in Supplementary Results). The parcellation was created for the left hemisphere and mirrored onto the right hemisphere using left-right surface registration to achieve left-right symmetry (Greve et al. 2013) (Figure S2). We note that the use of a random atlas does not allow for direct comparison of individual regions across species in terms of anatomical location or function, but that it does ensure that resulting connectivity maps include the same number of evenly spaced regions across species (van Wijk et al. 2010). Furthermore, the used implementation defined matching brain areas across the two hemispheres within each species, allowing for within-species comparisons of connectivity profiles (Mars et al. 2016) of spatially corresponding regions across the left and right hemispheres (see *Connectivity asymmetry*).

### Connectome reconstruction

DWI datasets were corrected for eddy current, motion, and susceptibility artifacts using FSL (Jenkinson et al. 2012), followed by deterministic fiber tracking and connectome reconstruction using CATO (v2.5, de Lange et al., submitted, www.dutchconnectomelab.nl/cato). Voxel-wise diffusion profiles were reconstructed using generalized *q*-sampling imaging (GQI) (Yeh et al. 2010) with a tensor model used in the absence of a complex fiber configuration (de Lange et al., submitted; Romme et al., 2017). Eight streamline seeds were started from each voxel in the white matter mask, with streamlines propagated along the best matching diffusion direction from voxel to voxel until one or more of the stop criteria was reached (exited the brain mask, made an angle of > 60°, fractional anisotropy of < 0.1). The set of cortical areas were used as network nodes, with connectivity between two network nodes defined as the number of streamlines reaching both representative cortical areas, resulting in a connectivity matrix describing the reconstructed corticocortical white matter pathways with the number of streamlines (NOS) of connections taken as a measure of connection strength.

### Connectivity asymmetry

Connectivity patterns of the left vs right hemispheres were compared by means of calculating the level of ‘connectivity asymmetry’, with the goal of capturing potential differences in connectivity between brain regions (Mars et al. 2016). For each region A in the left hemisphere we calculated how strongly region A was connected to other regions in the left hemisphere (i.e., the connectivity profile of region A). The same was performed for the spatial homolog A’ of region A in the right hemisphere. We then computed the difference between these two connectivity profiles by calculating the absolute difference in connection strength for each of the connections in the two profiles. These per-connection difference scores were then averaged to obtain a connectivity asymmetry score for region pair A-A’. The asymmetry score for this region pair thus reflects the degree to which connection strength differs along the connections of region A compared with its homolog region A’, with a value of 0 denoting identical (i.e. symmetrical) connectivity profiles of A and A’ and values >0 denoting higher levels of asymmetry between the two regions. To ensure scores reflected differences in connection strength, we focused on connections present for both regions A and A’. Finally, connectivity asymmetry scores were averaged across all left-right pairs of regions, resulting in a total connectivity asymmetry score for each dataset.

We took further normalization steps to ensure that the measure of connectivity asymmetry itself was unaffected by any potential cross-species differences in overall number of reconstructed connections or overall strength of these connections. Specifically, networks were controlled for differences in total number of reconstructed connections by setting network density (i.e. the total number of connections) equal across species (van Wijk et al. 2010; van den Heuvel et al. 2017) and controlled for global differences in total connectivity strength by resampling the weights to a normal distribution with identical mean and standard deviation (*M* = 1, *SD* = 0.2) for both hemispheres in each species (Hagmann et al. 2008; Honey et al. 2009) (for additional analysis based only on presence and absence of connections see Supplementary Results). The examined metric of connectivity asymmetry thus reflected relative differences in the distributions of connection weights between the left and the right hemispheres within each species, allowing comparison of the resulting normalized values across species.

### Statistical analysis

Associations between predictor and outcome variables were tested by means of phylogenetic generalized least squares regression (PGLS) (Pagel 1997; Symonds and Blomberg 2014), an extension of ordinary least squares regression that accounts for potential non-independence of the comparative data due to shared evolutionary history. PGLS was performed with the caper package in R (Orme et al. 2018) using a consensus phylogenetic tree obtained from the 10kTrees project version 3 (Arnold et al. 2010). A Brownian motion model of evolution (Felsenstein, 1985) was fitted, modeling phenotypes of species that share a recent common ancestor to be more similar than phenotypes of more distantly related species. The strength of the evolutionary signal was measured as the covariance in the residuals and captured by Pagel’s *λ*, with values varying between 0 (no phylogenetic signal in the residuals) and 1 (expected covariance under a Brownian motion model of evolution). PGLS analyses of volumetric and surface properties were conducted on log-transformed data such that the regression coefficients could be interpreted as scaling exponents. PGLS analyses of connectivity asymmetry were conducted on *z*-transformed data, yielding standardized regression coefficients.

### Data sharing and code accessibility

Raw MRI data are available from the sources listed in Table 1. Processed volumetric and connectivity data, as well as code for data analysis and figures are made available upon publication at github.com/ardesch/brainscaling. Raw and processed MRI data from the primate species included from the Primate Brain Bank are available at primatebrainbank.org/digital.

## Results

### Gray and white matter

We started by first quantifying the global scaling relationships between the gray matter, white matter, and cortical surface. Cerebral volume varied over 350-fold across the fourteen species, from 2.6 cm^3^ for the galago to 905 cm^3^ on average for the human subjects (Figure 2A). If larger brains were simply isometrically scaled or ‘blown-up’ versions of smaller brains, cortical surface area would scale to the 2/3 power of cerebral volume, resulting in an expected range in surface area of 350^2/3^ = 50-fold in our sample. Cortical surface area instead varied over 140-fold, between 11 cm^2^ for the galago and 1555 cm^2^ on average for the human subjects (Figure 2A). Confirming comparative observations of earlier MRI studies (Rilling and Insel 1999b; Mota et al. 2019), cortical surface area was found to outpace cerebral volume with a positive allometric scaling exponent of b = 0.85 (95% confidence interval (CI) = 0.82-0.88, adjusted R^2^ = 0.99, Pagel’s *λ* = 0, P < 2 × 10^-16^). This exponent of 0.85 exceeds an exponent of 2/3 for simple isometric scaling of a two-dimensional surface against a three-dimensional volume (Figure 2B), which indicates that larger brains have proportionally more cortical surface area than expected for their cerebral volume.

**Figure 2:**
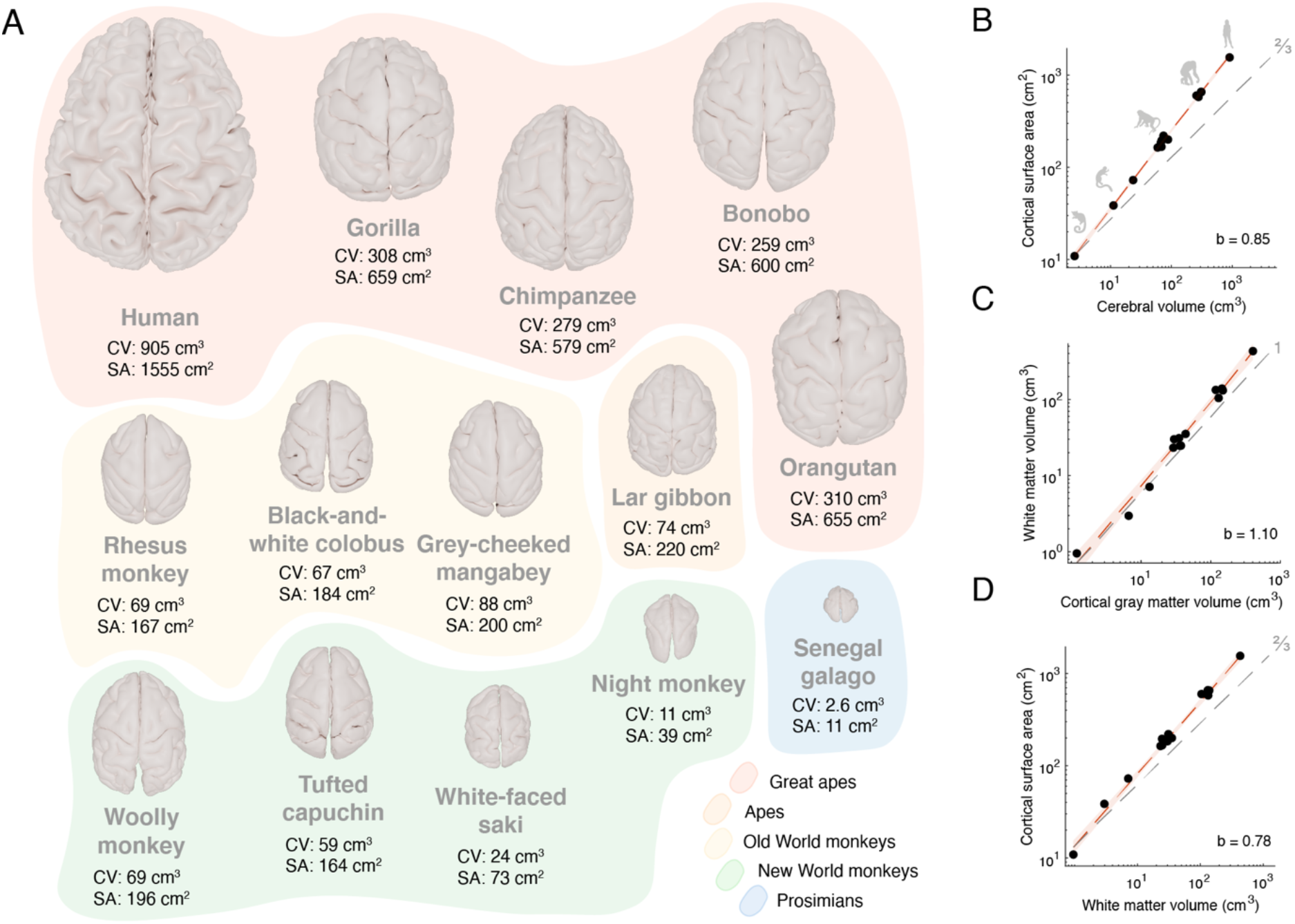
Scaling relationships of the cerebrum. A) Cortical surface reconstructions (to scale). Cerebral volume (CV) and cortical surface area (SA) were computed from the structural T1 and T2* MRI datasets. B) Scaling between cortical surface area and cerebral volume shows a strong positive allometric relationship. C) White matter volume scales with positive allometry on cortical gray matter volume. D) Cortical surface area scales with positive allometry on white matter volume. In plots B-D, the dotted gray line indicates isometric scaling and is annotated with the expected slope for isometric scaling between a surface and a volume (2/3), and between two volumes (1). 95% confidence bands are plotted in red to indicate positive allometry. Scaling formula: log(y) = b · log(x) + intercept.

White and gray matter volume further tended to scale with positive allometry with a scaling exponent of b = 1.10 (95% CI = 0.99-1.21, adjusted R^2^ = 0.97, Pagel’s *λ* = 0.54, P = 4.8 × 10^-11^), corresponding to an increase in the proportion of white matter to total cerebral volume from 37% in the galago to 39% in the capuchin monkey, 43% in the gorilla and 48% in humans (Figure 2C). Further examining cortical surface area across species showed that total area of the cortical mantle outpaces total cerebral white matter volume (b = 0.78, 95% CI 0.73-0.83, Pagel’s *λ* = 0.60, P = 4.2 × 10^-13^) (Figure 2D), indicating that while the proportion of total volume devoted to cerebral white matter volume is higher in larger brains, it may not be able to keep pace with the rapid cortical expansion in larger brains.

### Corpus callosum

Extending these previous MRI observations, we then continued by examining more specifically the scaling relationships between cortical surface and the corpus callosum (CC), the brain’s largest white matter bundle interconnecting regions across the two hemispheres (Figure 3A). We particularly wanted to examine whether an increase in the proportion of white matter is able to keep up with an increasing demand of interregional connectivity between the hemispheres in larger brains. Corpus callosum cross-sectional area was found to scale with negative allometry on cortical surface area (b = 0.88, 95% CI = 0.81-0.95, adjusted R^2^ = 0.98, Pagel’s *λ* = 0, P = 4.0 × 10^-12^, Figure 3B), indicating that the corpus callosum does not keep pace with cortical surface area. The space available for interhemispheric connectivity thus does not keep up with the larger cortical surface of bigger brains, with 1 cm^2^ of CC area available per 90 cm^2^ of cortical surface in the galago but only 1 cm^2^ of CC area per 132 cm^2^ of cortical surface in chimpanzees and only 1 cm^2^ of CC area per 211 cm^2^ of cortical surface in humans.

**Figure 3:**
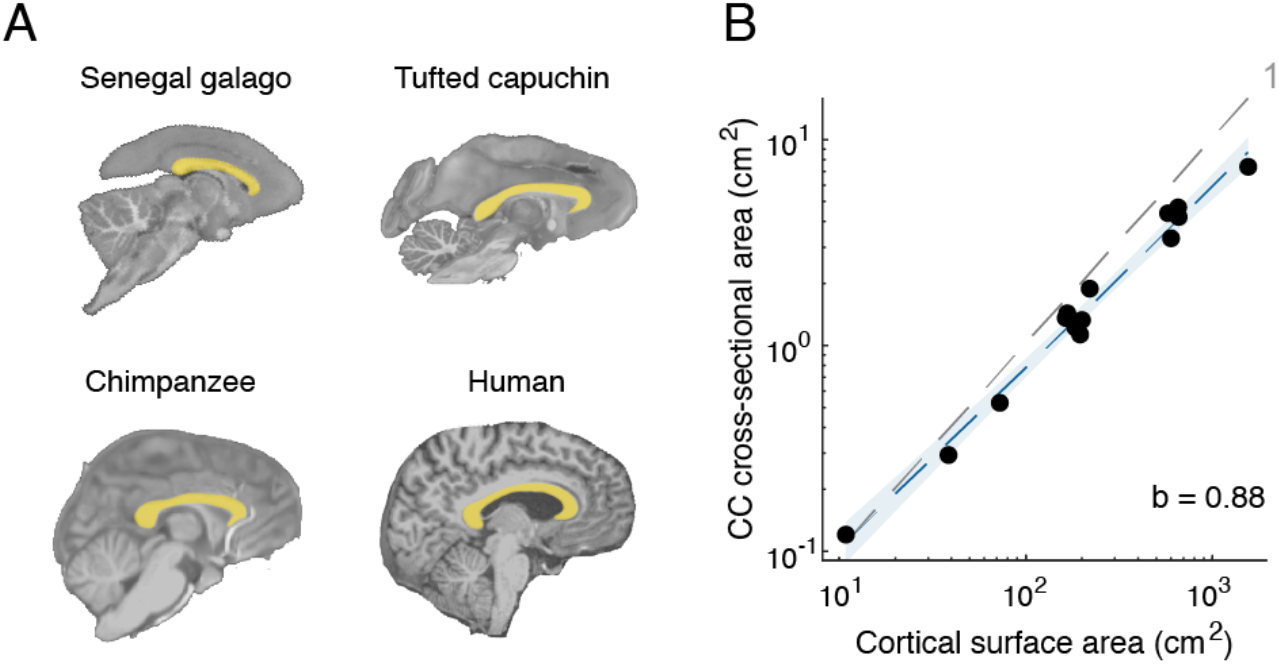
Scaling relationships of the corpus callosum. A) Segmentation of the crosssectional area of the corpus callosum (CC, yellow overlay) in a sagittal slice of the Senegal galago, tufted capuchin, chimpanzee, and human brain (not to scale). B) CC cross-sectional area scales with negative allometry on cortical surface area. The dotted gray line indicates the expected slope of 1 for isometric scaling between two surfaces. 95% confidence band is plotted in blue indicating negative allometry.

### Macroscale connectivity patterns

The findings of cortical surface area outpacing white matter volume and particularly the corpus callosum suggest a potential ‘bottleneck’ in interhemispheric connectivity in larger brains. As a third analysis, we therefore investigated the implications of brain scaling relationships on the network organization of brain connectivity. We calculated connectivity asymmetry, a normalized measure of connectivity differences between the left and the right hemispheres (Figure 4A, see also Materials and Methods). Connectivity asymmetry was found to be significantly higher in larger brains, indicative of more specialized brain network organization in larger-brained species (standardized β = 0.73, 95% CI = 0.27-1.18, adjusted R^2^ = 0.48, Pagel’s *λ* = 0, P = 5.0 × 10^-3^, Figure 4B). Connectivity asymmetry similarly correlated to cortical surface area (standardized β = 0.73, 95% CI = 0.28-1.18, adjusted R^2^ = 0.49, Pagel’s *λ* = 0, P = 4.5 × 10^-3^), indicating that connectivity patterns in the left and right hemispheres become increasingly asymmetrical with increasing brain volume and cortical surface area. These effects were controlled for variation in both absolute number and total strength of connections across species, validated across comparison to effects in brain networks with randomly permuted connection strengths (P < 2 × 10^-16^, 1,000 permutations, Figure 4B inset), and replicated using alternative measures of connection strength (Supplementary Results).

**Figure 4:**
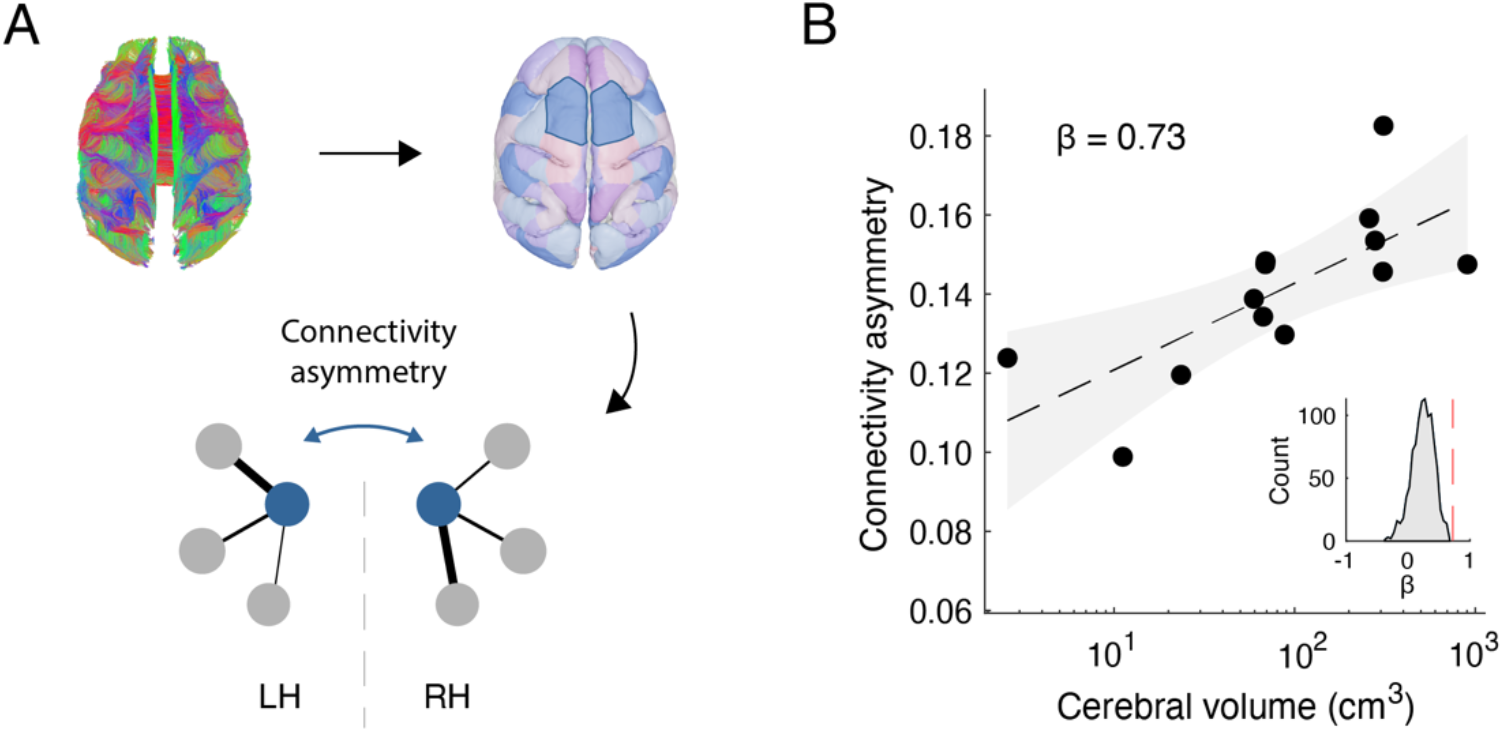
Scaling between cerebral volume and connectivity patterns. A) Fibers reconstructed from diffusion-weighted imaging (top left panel) were combined with a leftright symmetric cortical parcellation (top right panel) to compare asymmetry of connectivity patterns between the two hemispheres in each species (tufted capuchin shown as an example). The toy network in the bottom panel depicts two spatially homologous regions (blue nodes) with asymmetrical patterns of connectivity strength (difference in connection thickness between left and right). B) Mean connectivity asymmetry of spatially homologous regions in the left vs the right hemispheres, plotted against cerebral volume (95% confidence band plotted in gray). Inset: Null distribution of the association between connectivity profile asymmetry and cerebral volume after randomly shuffling the connection weights (1,000 permutations). The red line represents the observed regression coefficient as depicted in the main figure.

## Discussion

Our study provides new insights into scaling relationships of major brain components and the consequences for macroscale brain network circuitry in the human and non-human primate brain. We first validate long-standing evidence of post-mortem and comparative MRI studies that white matter volume is outpaced by a growing surface area of the cortical mantle in larger brains, resulting in relatively less space for connectivity to keep cortical areas equally connected in larger brains. Our findings further extend this by showing that a relative reduction in brain connectivity may result in a potential constraint on long-range connectivity in larger brains, prompting higher levels of lateralization of brain regions in the left and right hemispheres.

Positioning the current findings in a broader context of global brain scaling relationships, the observed relationships between gray matter volume, white matter volume, and cortical surface area corroborate previous comparative studies (Hofman 1989; Rilling and Insel 1999b; Rilling and Insel 1999a; Mota et al. 2019). Our comparative data across a 350-fold range in brain size shows that the proportion of total cerebral volume devoted to white matter connectivity tends to increase exponentially from smaller to larger brains, from around one third in smaller primates to almost half the volume (48%) in the human brain. Indeed, the exponent of 1.10 between white matter and gray matter in our data is in line with previous estimates of 1.12 (Rilling and Insel 1999a) and 1.14 (Mota et al. 2019), underlining the robustness of these allometric relationships across different datasets and methodologies.

Higher proportions of white matter have been proposed to help connect the expanding cortex in larger brains, which requires disproportionately more white matter to remain connected (Deacon 1990; Ringo 1991; Hofman 2012). Despite this growing white matter proportion, our findings show that cortical surface area grows even faster and outpaces white matter volume and the corpus callosum, closely in line with earlier findings (Rilling and Insel 1999b). The observed negative scaling exponent of 0.88 shows that a 2-fold increase in cortical surface area corresponds to a growth of corpus callosum cross-sectional area of 1.84-fold (= 2^0.88^), indicating that there is less and less space available for callosal connectivity per unit of cortical surface area in larger brains and pointing to the emergence of a long-distance connectivity bottleneck in larger brains.

A bottleneck in long-range connectivity and a related increase in asymmetry in brain network organization in larger brains have been suggested as an important source for functional lateralization in large-brained primates (Barrett 2012; Rilling 2014; Phillips et al. 2015). Brain lateralization has been suggested to be an important catalyst for the evolution of specialized and advanced cognitive functions, such as the emergence of complex language and social intelligence in humans (Ringo et al. 1994; Gazzaniga 2000; Roth and Dicke 2005; Rilling et al. 2008; Barrett 2012). The balance between local, more specialized connectivity and high degrees of global interconnectivity might present an important trade-off in brain organization, as shown by a large cross-species study across a broad range of animal orders (Assaf et al. 2020). Indeed, lower levels of global connectivity together with higher levels of lateralization could lead to reduced network redundancy. This has been suggested to render the human brain potentially more vulnerable to brain damage such as stroke (Bartolomeo and Thiebaut de Schotten 2016) and neurodevelopmental disorders (Ribolsi et al. 2009; Bishop 2013; Xie et al. 2018). The corpus callosum, one of the largest white matter bundles of the brain and the main bridge between the hemispheres, is involved in a wide range of both neurological and psychiatric illnesses (de Lange et al. 2019), including amyotrophic lateral sclerosis (Filippini et al. 2010), schizophrenia (Prendergast et al. 2018), and bipolar disorder (Francis et al. 2016).

The observed brain scaling relationships suggest that the human brain is not particularly different or ‘unique’ compared to a general primate trend: Although the human brain is larger than expected for a typical primate of the same body size (Rilling 2014), the contribution of the major tissues to its overall size is predictable (see also Azevedo et al. 2009; Miller et al. 2019). Our comparative connectome analyses now similarly suggest that aspects of brain network topology could also be direct consequences of brain scaling principles. Indeed, the current findings support the notion that a high degree of network specialization in the human brain is inherently linked to higher wiring costs associated with an increasing proportion of cortical surface area in larger brains (van den Heuvel et al. 2016).

A number of methodological factors need to be taken into account when interpreting our findings. The scaling relationships found are fitted on a representative sample across a range of monkeys and apes but do not necessarily generalize to all species (Jyothilakshmi et al. 2020); positive allometry between white and gray matter scaling is a conserved feature in primates, but absent in artiodactyls (hoofed animals including giraffes and deer) (Mota et al. 2019). A second point to consider is that our current analyses are limited to the cerebrum and corticocortical connections. The cerebral cortex has been an important structure of interest in many comparative studies, but the cerebellum has also undergone rapid changes in the evolutionary branch leading up to apes (Miller et al. 2019; Smaers and Vanier 2019). The cerebellum was unfortunately not preserved in some of the post-mortem samples we examined and we therefore had to exclude this structure from our analyses. Third, it needs to be mentioned that while diffusion-weighted imaging is one of only a few methods available to obtain information on white matter connections both in vivo and post-mortem, and while it tends to show reasonable agreement with more direct and invasive methods such as tract tracing (van den Heuvel et al. 2015; Delettre et al. 2019), diffusion-weighted imaging is well known to suffer from a range of methodological limitations. These limitations result in both false positive and false negative fiber reconstructions (Thomas et al. 2014; Maier-Hein et al. 2017), which in turn have their effect on network reconstruction and analysis (de Reus and van den Heuvel 2013; Zalesky et al. 2016). We aimed to minimize the potential impact of false positives by focusing on connections present in both the left and right hemispheres in our weighted network analyses, and by averaging connectivity profile differences across regions. Nevertheless, further investigations using methods at the meso- and microscale will be necessary to specify which types of cortical areas, fiber bundles, and neurons may underlie the connectivity scaling patterns observed on the macroscale.

## Acknowledgements

We thank John Allman for sharing the post-mortem great ape data used in this study. This work was supported by the Netherlands Organization for Scientific Research (NWO) (grant numbers 452-16-015, ALWOP.179 to M.P.v.d.H., grant number 452-13-015 to R.B.M.); the Biotechnology and Biological Sciences Research Council (grant number BB/N019814/1 to R. B.M.); the Wellcome Trust (203139/Z/16/Z); the National Institutes of Health (grant number P01 AG026423 to T.M.P. and J.K.R.), the National Center for Research Resources (grant number P51RR165 to T.M.P. and J.K.R. (superseded by the Office of Research Infrastructure Programs/OD P51OD11132)), the National Chimpanzee Brain Resource (grant number R24NS092988 to T.M.P. and J.K.R.), and Amsterdam Neuroscience (alliance grant to S. C.d.L.).

