## Supplementary Material for "Scaling principles of white matter brain connectivity"

### Supplementary Results

**Binary connectivity profile divergence.** Connectivity profile divergence in the main text was computed based on weighted connectivity, taking into account the strength of each connection. This analysis was based on connections present in both hemispheres, resulting in their binary connectivity patterns to be identical. As an alternative, we here computed connectivity profile divergence based on binary connectivity (describing the presence/absence of connections without information on connection strength). Region-wise overlap was calculated by computing for each homologous region pair the proportion of overlapping connections between the left and right hemispheres relative to the total number of connections of that region in either hemisphere, varying between 0 (no overlap) and 1 (complete overlap). The mean divergence in binary connectivity profiles was then obtained by averaging the region-wise overlap for each dataset and subtracting this number from 1. Correlating binary connectivity profile divergence with cerebral volume showed a trend for a positive association (standardized $\beta = 0.49$, 95% CI = -0.085-1.1, adjusted $R^2 = 0.17$, Pagel's $\lambda = 0$, $P = 0.09$). The higher standardized regression coefficient (0.73 vs 0.49) and stronger statistical significance ($P = 0.005$ vs $P = 0.09$) of the main weighted analysis compared with the binary analysis suggests that the positive association between connectivity divergence and brain size is most strongly driven by differences in the strength of connections between the left and right hemispheres, with potentially a more modest contribution from differences in binary connectivity patterns.

**Alternative connection weights.** Diffusion-weighted imaging protocols yield multiple metrics for each reconstructed fiber, each highlighting different aspects of the reconstructed connectivity. The main analysis of connectivity profile divergence was based on the number of streamlines (NOS), a metric that has been related to connection strength (e.g. Ardesch et al. 2019; Assaf et al. 2020). Here we repeat these analyses using fractional anisotropy (FA), another frequently used metric that is thought to relate to fiber microstructure with contributions from myelination and axonal structure (Alba-Ferrara and de Erausquin 2013). FA-based connectivity profile divergence showed a positive association with cerebral volume ($\beta = 0.60$, 95% CI = 0.074-1.13, adjusted $R^2 = 0.30$, Pagel's $\lambda = 0$, $P = 0.03$), exceeding effects of the null condition with randomly permuted connection weights ($P = 5.5 \times 10^{-3}$, 1,000 permutations). The FA-based divergence scores thus extend the effects reported in the main text by showing a similar association with brain size as the divergence scores based on NOS-based connection strength.

**Alternative cortical atlases.** Network analyses can be sensitive to the number of nodes (cortical areas) used (van Wijk et al. 2010). We chose an atlas containing 50 areas per hemisphere in the main text as a reasonable number of areas to use across species (similar to other often-used atlases such as the Desikan-Killiany atlas (Desikan et al. 2006) and the Yeo 17 functional network atlas (Yeo et al. 2011). We here repeated the analyses correlating cerebral volume to connectivity profile divergence using atlases of 25 and 100 areas per hemisphere, respectively. Using 25 areas per hemisphere showed a trend for a positive correlation between cerebral volume and connectivity profile divergence ($\beta = 0.49$, 95% CI = -0.087-1.06, adjusted $R^2 = 0.17$, Pagel's $\lambda = 0$, $P = 0.09$) with this positive correlation being significantly higher than the null model of randomly shuffled connection weights ($P = 0.01$, 1,000 permutations). Using 100 areas per hemisphere did not show a significant positive correlation ($\beta = 0.31$, 95% CI = -0.31-0.94, adjusted $R^2 = 0.02$, Pagel's $\lambda = 0$, $P = 0.30$), although this effect was higher than the correlation obtained in the null model ($P = 8.6 \times 10^{-3}$, 1,000 permutations). It is possible that this high number of 100 cortical areas per hemisphere goes beyond the number of cortical areas that may biologically be present, especially in the smaller brains, resulting in a connectivity network with nodes that are not sufficiently distinct from each other. Taken together, these analyses using different cortical atlases are in agreement with the main results in terms of the direction of the correlation, but the strength of the association seems to be at least partly dependent on the number of areas included in the atlas.

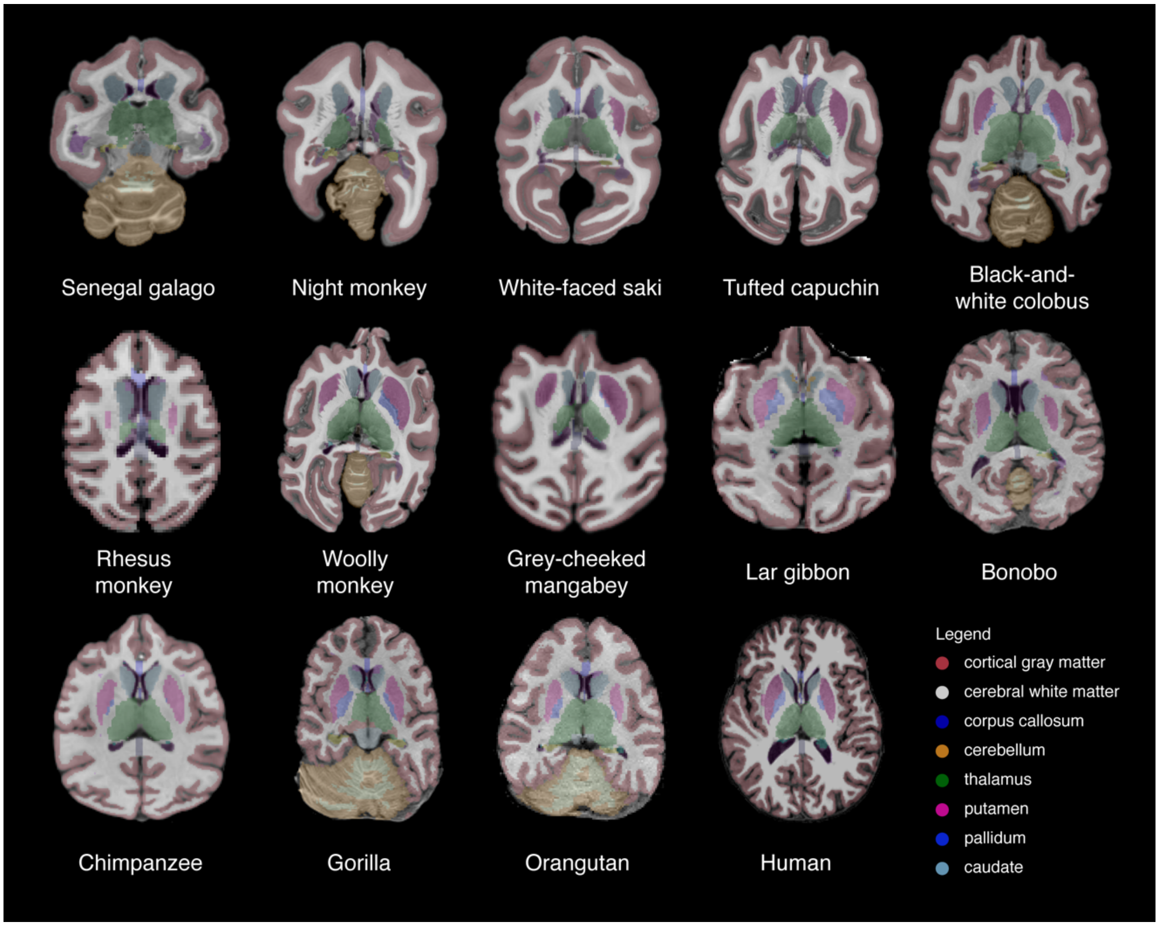


**Supplementary Figure 1.** Segmentation into major brain tissue classes for each species included in the study. The segmentation is overlaid on a bias field-corrected structural volume and depicted in a transverse slice through the thalamus. An exemplary subject was chosen for species with multiple samples/subjects. Brains are visualized at approximately the same size for comparison.

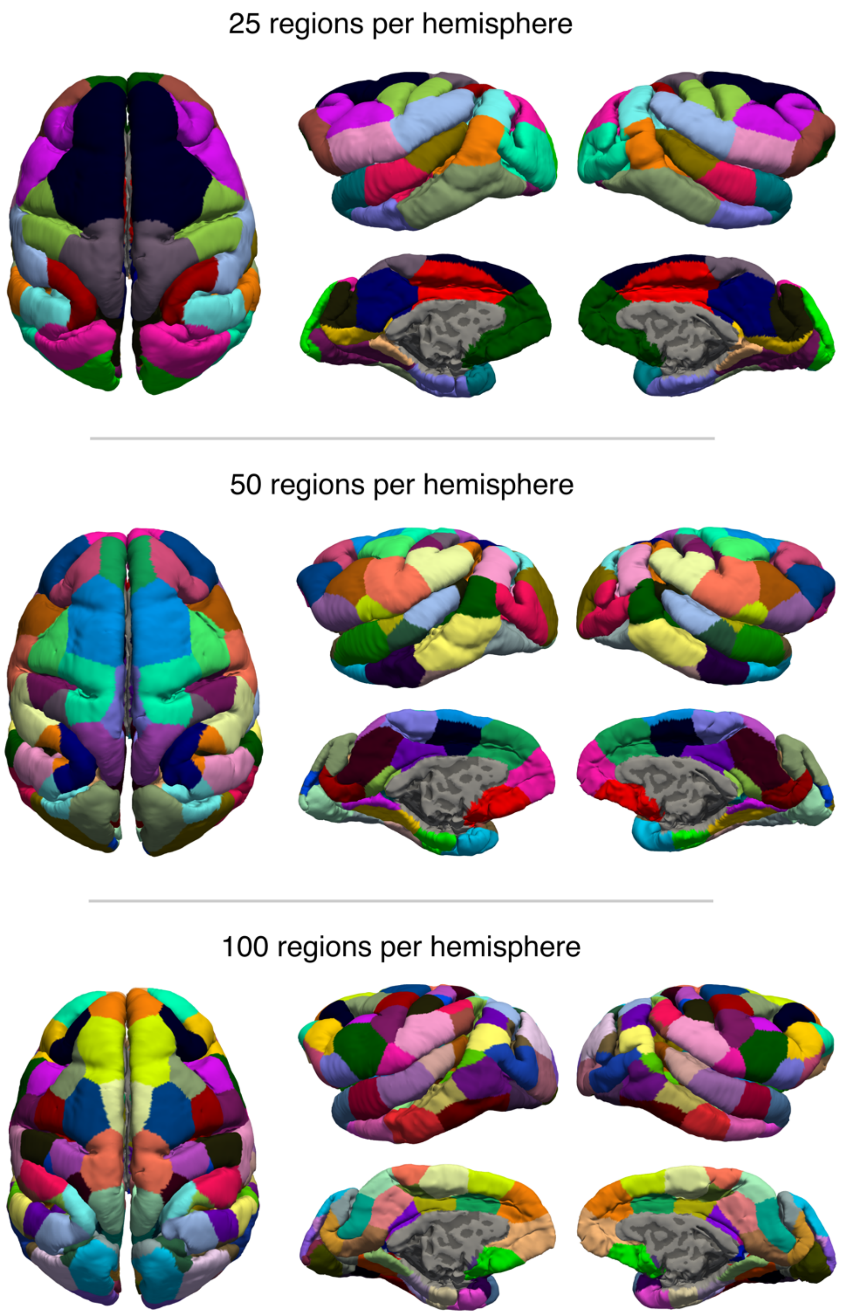


**Supplementary Figure 2.** Random left-right symmetrical cortical parcellation. Top-view (left), and lateral/medial views of the left hemisphere (middle) and right hemisphere (right)

are shown for the random atlases containing 25, 50, and 100 regions per hemisphere, respectively. The atlas is displayed on the cortical surface of the tufted capuchin.
